# Synergic Combination of Stimulated Emission Depletion Microscopy with Image Scanning Microscopy to Reduce Light Dosage

**DOI:** 10.1101/741389

**Authors:** Giorgio Tortarolo, Marco Castello, Sami Koho, Giuseppe Vicidomini

## Abstract

Stimulated emission depletion (STED) microscopy is one of the most influential nanoscopy techniques; by increasing the STED beam intensity, it theoretically improves the spatial resolution to any desired value. However, the higher is the dose of stimulating photons, the stronger are the photo-bleaching and photo-toxicity effects, which potentially compromise live-cell and long-term imaging. For this reason the scientific community is looking for strategies to reduce the STED beam intensity needed to achieve a target resolution. Here, we show how the combination of STED microscopy with image scanning microscopy (ISM) meets this request. In particular, we introduce a new STED-ISM architecture – based on our recent single-photon-avalanche-diode (SPAD) detector array – which allows covering the near-diffraction limit resolution range with reduced STED beam intensity. We demonstrate this ability both with simulated data and in live-cell experiments. Because of (i) the minimal changes in the optical architecture of the typical point-scanning STED microscope; (ii) the parameter-free, robust and real-time pixel-reassignment method to obtain the STED-ISM image; (iii) the compatibility with all the recent progresses in STED microscopy, we envisage a natural and rapid upgrade of any STED microscope to the proposed STED-ISM architecture.

---

The vast family of fluorescence optical microscopy techniques stands as an invaluable tool for addressing various biological questions, allowing to observe selectively and often for an extended time a given mechanism of interest, also directly in living cells or tissues. The so-called super-resolution techniques have overcome the long-standing spatial resolution limitation [1, 2, 3] by avoiding the simultaneous signaling of molecules closer than the diffraction limit. Stimulated emission depletion microscopy (STED) [4, 5], one of the first of such techniques, relies on an additional laser beam (the STED beam) whose focal intensity distribution is shaped as a doughnut to confine fluorescent only in the very centre of a conventional Gaussian excitation region. Notably, the higher the STED beam intensity, the smaller the effective fluorescent region, and ultimately the better the spatial resolution. Whereas STED microscopy attains single-digit nanometer resolution for some particularly tailored implementations [6], its practical performance is hindered by the non-negligible chance of photo-damaging the sample with the high-intensity STED beam. Despite several strategies to mitigate this problem have been proposed, such as time-resolved STED [7, 8] microscopy, adaptive-smart scanning schemes [9, 10, 11], and far-red illumination [12], it still holds as a significant concern when one performs a live-cell or long-term measurements. Therefore, the scientific community is still looking for strategies to reduce the STED beam intensity needed to achieve a target resolution.

Image scanning microscopy (ISM) [13, 14] stands as a promising candidate to meet this need: this technique is known to boost the spatial optical resolution and the signal-to-noise Ratio (SNR) of confocal microscopy, and can potentially provide a similar enhancement in other microscopy approaches, e.g. two-photon excitation microscopy or STED microscopy. In the context of laser point-scanning microscopy, we recently proposed a simple ISM implementation that relies on the usage of a single-photon-avalanche-diode (SPAD) detector array, instead of the typical single-element detector [15]. A single scan results consequently in a set of “similar” images, the so called scanned images, that are in principle only shifted with respect to the reference image generated by the central detector of the array. The simple pixel-reassignment (PR) operation, i.e., shifting back and summing all the images up, grants the spatial resolution and the SNR boost of the final ISM reconstruction. Here we show how this enhancement, in the context of STED microscopy, allows to reduce the STED power needed to achieve a target resolution. Notably, the proposed combination of STED and image-scanning microscopy (i) requires only minimal changes in the conventional STED microscopy architecture: an extra magnification on the image plane and the introduction of the SPAD array instead of the single-element detector (Supp. Fig. S1), (ii) is fully compatible with all the approaches for photo-damage reduction mentioned above, and (iii) preserves all the other functions of STED microscopy, such as multi-color, three-dimensional, and fast imaging.

The correct amount of shift – the shift vector – imposed to a given scanned image for the PR reconstruction method is key for a successful resolution and SNR enhancement. In first approximation, and in absence of stimulating photons, i.e., no STED imaging, the shift-vectors are theoretically related to the physical geometry of the detector array, and their modules equal half of the distance between the given sensitive element of the detector array and the central one, projected onto the sample plane [16]; however, the shift-vectors are practically influenced by the spectral properties of the fluorophores [17], and by any sample- or system-induced aberration or misalignment of the optical system [15]. Additionally, in the case of STED microscopy, the shift-vectors show a strong dependence on the STED beam intensity: the higher the photon dosage, the smaller the effective fluorescent spot, and the shorter the shift-vectors. In essence, the overall PSF of a STED microscope in the case of high STED intensities depends mainly on the excitation PSF, whilst the influence of the emission PSF becomes negligible, and thus the scanned images differ each other in SNR but not anymore in position. With this insight, we first show – through simulations – that the pixel reassignment operation is beneficial for STED microscopy for a given range of STED intensities; more precisely, the spatial resolution and the SNR of the STED-ISM reconstruction are enhanced with respect to the raw STED counterpart, for STED intensities below a given threshold (Fig. 1a, Supp. Fig. S2). Increasing the STED beam power more leads to very short shift-vectors and ultimately to negligible benefits from the pixel reassignment operation.

**Figure 1:**
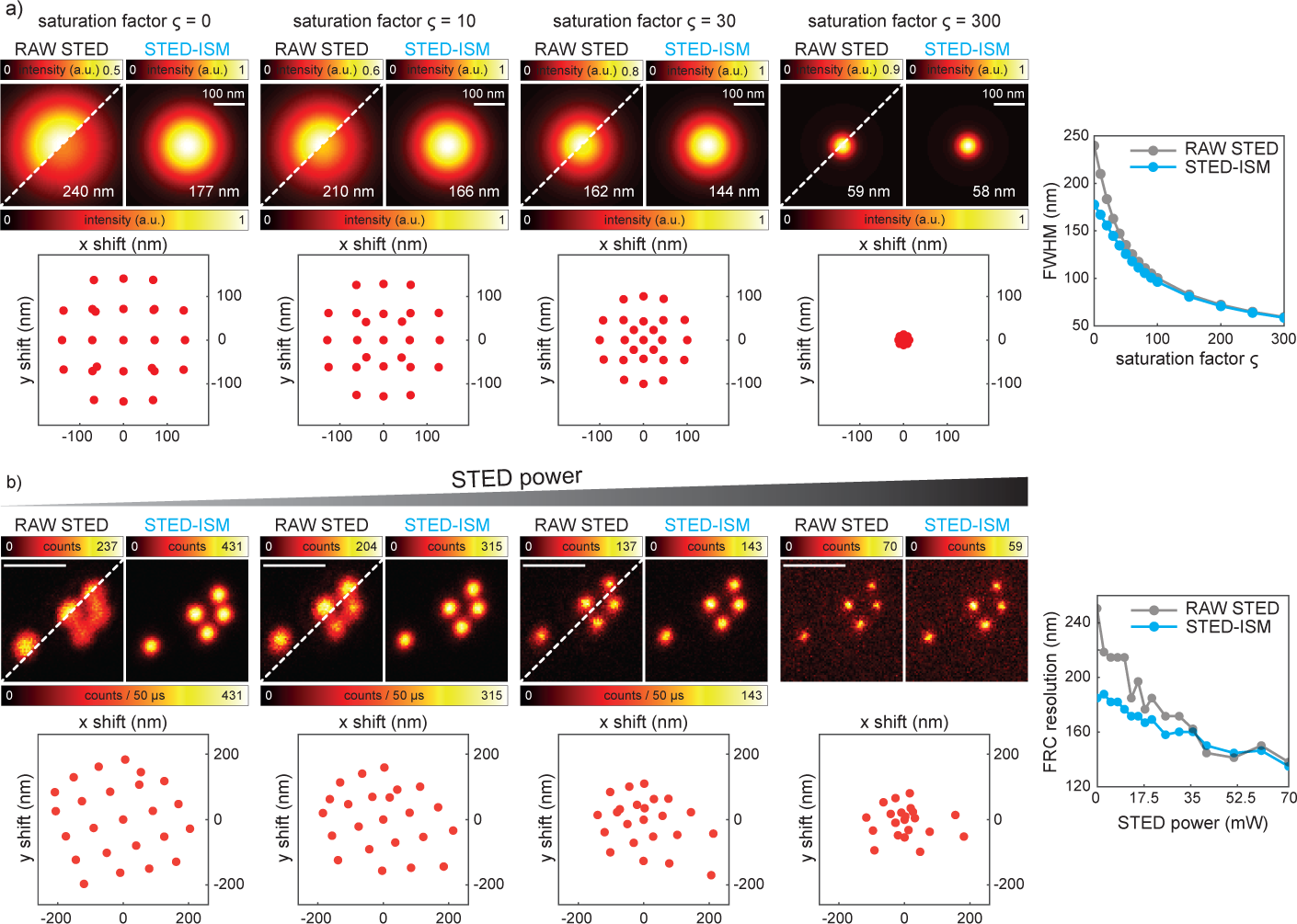
STED-ISM simulation and proof-of-principle. **a**, Simulation of the STED-ISM performances with increasing STED saturation factor *ζ* (from left to right). The STED-ISM Point Spread Function (PSF) is narrower and higher than the raw STED counterpart (top), resulting in a boost of spatial resolution and SNR; full-width-half-maximum analysis values (FWHM, right) are also reported on the images. As *ζ* increases and the effective excitation PSF shrinks, the images recorded by the 25 sensitive elements of the SPAD array are less and less shifted with respect to the one corresponding to the central element; the retrieved shift values decrease accordingly (bottom; for readability, the corresponding SPAD elements are not reported), leading ultimately to negligible advantages of the STED-ISM result. Excitation, emission and STED wavelengths: 640, 670 and 775 nm respectively; STED repetition rate: 80 MHz; STED pulse length: 200 ps; SPAD array semi-length: 1.4 Airy Units. **b**, STED-ISM imaging of 20 nm sized fluorescent beads with increasing STED power. The visual inspection of the details (top) suggests the superior performance of STED-ISM, confirmed by the FRC analysis (right). Dwell time: 50 *µ*s; pixel size: 20 nm; STED powers: 0, 13, 50, 160 mW; details format: 150 × 150 pixels.

We present various experiments to validate our STED-ISM implementation with a SPAD array. We started imaging 20 nm diameter fluorescent beads with increasing STED intensities (Fig. 1b): as expected from simulations, the STED-ISM image shows better SNR when compared to the raw STED counterpart, obtained summing all the 25 channels up. The spatial resolution, here measured using Fourier Ring Correlation (FRC)[18], is also enhanced within a given STED intensity range. We confirmed these results by performing STED-ISM measurements of a sample of tubulin-labelled fixed Hela cells (Supp. Fig. S3). It is important to note here that the successful STED-ISM reconstruction relies heavily on a blind, parameter-free phase-correlation algorithm that we developed previously [19, 15] (Supp. Note 1), capable of retrieving the pixel reassignment shift-vectors directly from the scanned images. Although in general this strategy compensates for eventual aberrations and misalignments of the optical system, in the context of STED-ISM it also accommodates implicitly the strong dependency of the shift-vectors on the STED intensity. Conversely, other all-optical ISM implementations – both single-spot [17, 20, 21] or eventually multi-spots [22, 23] architectures – should rely on proper modelling or prior calibration of the STED microscope’s effective PSF, as a function of the STED beam intensity and of the general experimental conditions.

To investigate the concrete advantages of STED-ISM over STED microscopy, we explored the conditions for which one is usually concerned about the maximum amount of stimulating photons delivered to the sample: live cell imaging (Fig. 2, Supp. Fig. S4). Also in this case, we report a SNR and spatial resolution enhancement of the resulting STED-ISM images with respect to the raw STED counterparts. The result is further improved applying our multi-image deconvolution algorithm [24, 15], here completely parameter-free thanks to the PSFs estimation *via* FRC analysis [25]. Moreover, we were able to perform extended STED-ISM time lapses of live Hela cells without inducing any noticeable photo-bleaching effect, given the reduced STED intensity necessary to obtain the target resolution (Supp. Fig. S5, Supp. Video 1, 2, 3, 4, 5, 6).

**Figure 2:**
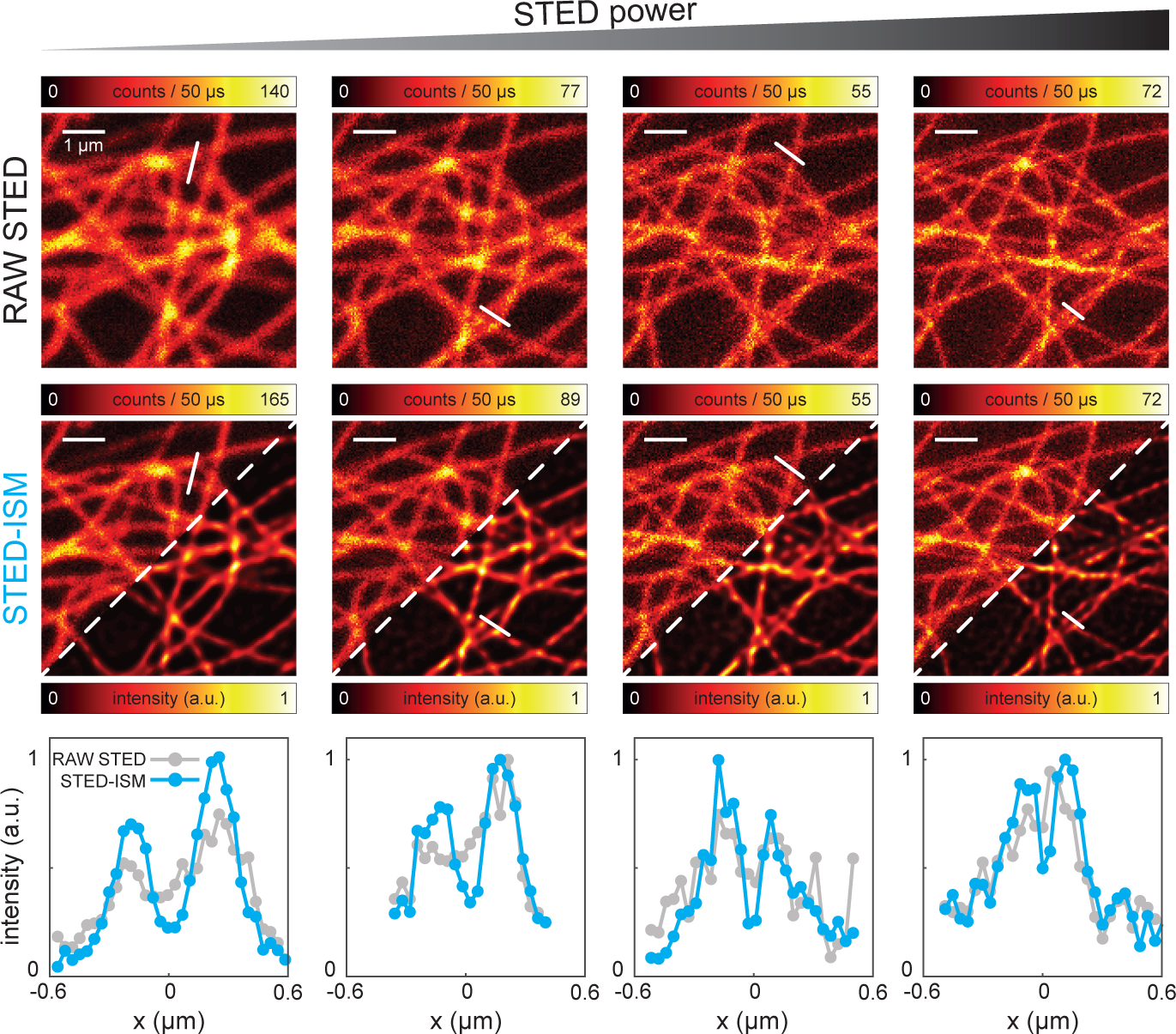
STED-ISM imaging of living cells. Raw STED (top), STED-ISM and the multi-image deconvolution result STED-ISM^+^ (center) details of SIR tubulin labelled living Hela cells with increasing STED powers (from left to right). The line profiles (bottom) proof the superior performances of STED-ISM with respect of the raw STED counterpart. Dwell time: 50 *µ*s; pixel size: 40 nm; STED powers: 0, 10, 20, 45 mW; details format: 150 × 150 pixels.

To conclude, we have introduced here for the first time a reliable and straight-forward combination of STED and ISM, thanks to our SPAD array-based ISM platform. Our approach allows to cover the near-diffraction limit resolution range (*λ/*(2 · *NA*) - 100 nm) with reduced STED beam intensity, mitigating the chances of photo-damaging the sample. The STED-ISM image shows enhanced SNR and resolution when compared to the raw STED counterpart, although the improvements obtained *via* the pixel reassignment approach become negligible when targeting resolutions far below the diffraction limit. The adoption of a parallel detection architecture clearly improves the dynamics - i.e., the maximum overall count rate - of the system [15, 26]; however, this benefit may not be relevant in the context of STED microscopy, considering its reduced fluorescence volume and thus photon flux when compared to confocal modality. Conversely, the photon detection efficiency (PDE) of the SPAD array detector is crucial. An interesting solution to increase the PDE relies on the adoption of the Bipolar-CMOS-DMOS (BCD) technology [27]; background-corrected deconvolution algorithms, coupled with one sole detector characterisation, would account for the probable consequent increase in dark noise.

Moreover, thanks to the single-photon timing nature of our SPAD array, a future synergy of ISM and time-resolved STED [28] would potentially allow for a computationally efficient way to recombine photons originated from different regions of the effective STED fluorescent volume. In conclusion, this work proves the benefits of ISM for STED microscopy, and marks an important step toward the diffusion of the SPAD array as a crucial upgrade for any point-scanning microscope.

## Acknowledgements

This work was partially supported by the Compagnia di San Paolo (ROL 20641 to G.T. and G.V.). We thank Prof. Alberto Tosi, Dr. Mauro Buttafava, and Federica Villa – from Politecnico di Milano – for the joint development of the SPAD array detector and for the continuous support in its optimisation; Prof. Alberto Diaspro, Prof. Colin J.R. Sheppard, Dr. Paolo Bianchini and Dr. Simonluca Piazza – from Istituto Italiano di Tecnologia – for useful discussions; Elena Tcarenkova – from University of Turku – for helpful discussions in the early stages of this project. Dr. Luca Lanzanó, Dr. Michele Oneto, and Simone Pelicci – from Istituto Italiano di Tecnologia – for support in sample preparation.

## Author contributions

G.T., M.C., and G.V. conceived the idea. G.V. supervised the project. G.T., M.C, S.K., and G.V. designed and implemented the custom STED-ISM system and developed the analysis and control software. G.T performed the experiments. G.T., and G.V. analyzed the data with support from all other authors. G.T., and G.V. wrote the manuscript with input from all other authors.

## ONLINE METHODS

### Custom setup

For this work, we updated the ISM setup described previously [1] with a STED line. Briefly, the excitation beam was provided by a triggerable pulsed (∼ 80 ps pulse-width) diode lasers (LDH-D-C-640, Picoquant) emitting at 640 nm. The STED beam was provided by a femtosecond mode-locked Ti:Sapphire laser (Chameleon Ultra II, Coherent) running at 775 nm. We coupled the laser beam into a 100 m long polarization maintaining fiber (PMF). Before injection into the PMF, the beam passed through two 20 cm long glass rods to temporally stretch the pulse-width to few picoseconds in order to avoid unwanted nonlinear effects and damages during the fiber coupling. We used a half-wave plate (HWP) to adjust beam polarization parallel to the fast axis of the PMF. The combination of glass rods and PMF stretched the pulses of the STED beam to ≈ 200 ps. We controlled the power of the STED and excitation beams thanks to two acoustic optical modulators (AOM, MCQ80-A1,5-IR and MT80-A1-VIS, respectively, AAopto-electronic). The Ti:Sapphire laser (master) runs at 80 MHz and provides an electronic reference signal which we used to synchronize electronically the excitation laser diode (slave). We used a picosecond electronic delayer (Picosecond Delayer, Micro Photon Devices) to temporally align the excitation pulses with respect to the depletion pulses. The STED beam emerging from the PMF is collimated, filtered in polarization by a rotating GlanThompson polarizing prism and phase-engineered though a polymeric mask imprinting 02*π* helical phase-ramps (VPP-1a, RPC Photonics). We rotated a quarter-wave plate and a half-wave-plate to obtain circular polarization of the STED beam at the back-aperture of the objective lens. We co-aligned the excitation and STED beam using two dichroic mirrors (T750SPXRXT and H643LPXR, AHF Analysentechnik). After combination, the excitation and STED beams were deflected by two galvanometric scanning mirrors (6215HM40B, CT Cambridge Technology) and directed toward the objective lens (CFI Plan Apo VC 60 ×, NA, Oil, Nikon) by the same set of scan and tube lenses used in a commercial scanning microscope (Confocal C2, Nikon). The fluorescence light was collected by the same objective lens, descanned, and passed through the multiband dichroic mirror as well as through a fluorescence band pass filter (685/70 nm, AHF Analysentechnik). A 300 mm aspheric lens (Thorlabs) focuses the fluorescence light into the pinhole plane generating a conjugated image plane with a magnification of 300×. For ISM measurements the pinhole is maintained completely open. A telescope system, built using two aspheric lenses of 100 mm and 150 mm focal length (Thorlabs), conjugates the SPAD array with the pinhole and provides an extra magnification factor. The final magnification on the SPAD array plane is 450×, thus the size of the SPAD array projected on the specimen is ∼ 1.4 A.U. (at the far-red emission wavelength, i.e. 650 nm). Every photon detected by any of the 25 elements of the SPAD array generates a TTL signal that is delivered through the dedicated channel (one channel for each sensitive element of the detector) to an FPGA-based data-acquisition card (NI USB-7856R from National Instruments), which is controlled by our own data acquisition/visualization/processing software, carma. The software-package carma also controls the entire microscope devices needed during the image acquisition, such as the galvanometric mirrors, the axial piezo stage, and the acousto-optic modulators (AOMs), and visualizes the data. In particular, carma synchronizes the galvanometric mirrors with the photon detection to distribute photons between the different pixels of the images. All power values reported for this setup refer to the power measured before the couple of galvanometric mirrors.

### Sample preparation

We demonstrated the enhancement in spatial resolution obtained with our STED-ISM approach on two-dimensional (2D) imaging of fluorescent beads and tubulin filaments. *Fluorescent beads.* In this study we used a commercial sample of ATTO647N fluorescent beads with a diameter of 23 nm (Gatta-BeadsR, GattaQuant). *Tubulin filaments imaging in fixed cells.* Human HeLa cells were fixed with ice methanol, 20 minutes at -20 °C and then washed three times for 15 minutes in PBS. After 1 hour at room temperature, the cells were treated in a solution of 3% bovine serum albumin (BSA) and 0.1% Triton in PBS (blocking buffer). The cells were then incubated with the monoclonal mouse anti-*α*-tubulin antiserum (Sigma Aldrich) diluted in a blocking buffer (1:800) for 1 hour at room temperature. The *α*-tubulin antibody was revealed using Abberior STAR Red goat anti-mouse (Abberior) for the custom microscope or Alexa 546 goat anti-mouse (Sigma Aldrich) for the Nikon-based microscope. The cells were rinsed three times in PBS for 5 minutes. *SiR-Tubulin in live-cells.* To label tubulin proteins, Human HeLa cell were incubated with SiR-tubululin kit (Spirochrome) diluted in LICS at a concentration of 1 *µ*M for 30 minutes at 37 °C and immediately after imaged at the microscope.

### Image reconstruction and analysis

To reconstruct the high-resolution ISM image we used either a simple pixel-reassignment method or a multi-image algorithm, which is fully described in Castello et al. [1]. Here, we briefly resume the two methods.

The pixel-reassignment (PR) method consists of (i) shifting each scanned image (*i, j*) of a shift-vector **s**_*i,j*_; (ii) adding up all the shifted images. In this work the shift-vectors are directly estimated from the scanned images, without the need of any input from the user. In particular, we use a phase correlation approach [2, 3] able to automatically take into the consideration the geometry of the detector array and the magnification of the microscope system, able to compensate for distortions (mis-alignments and aberrations) of the system which may arise during imaging. Very importantly in the context of this work, this automatic estimation of the shift-vectors accounts for the saturation level of the STED experiment, i.e. how much the effective fluorescence volume is shrunk, without the need of laborious calibration procedures. The multi-image deconvolution is routinely used when it is necessary to fuse different microscopy images of the same sample, but characterised by different point-spread-functions (PSFs) [4, 5, 6, 7, 8]. Here, we used the multi-image generalization of the well-known Richardson-Lucy algorithm:

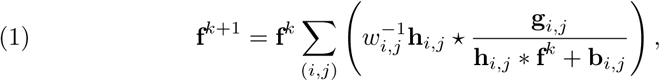

where **h**_*i,j*_ is the PSF linked to the element (*i, j*) of the SPAD array, **g**_*i,j*_ the series of scanned images, **f**^*k*^ the reconstructed image at the iteration *k, w*_*i,j*_ = (0, .., 1] a scaling factor, that takes into account the different SNR of the scanned images. With respect to the multi-image deconvolution algorithm introduced in Castello et al. [1], we introduce the term **b**_*i,j*_, which is the expected background associated to the scanned images **g**_*i,j*_. This terms can be used to introduce into the algorithm information such as the anti-Stokes emission background induced by the STED beam [9, 10], and/or the dark noise introduced by the SPAD array detector. While the expected anti-Stokes emission background depends on many different parameters and may require dedicated acquisition schemes to estimate, the expected dark-noise can be easily measured by registering the signal from the SPAD in “dark” conditions, i.e, in absence of any source of light. In this work, the Stokes-shift background is negligible and the estimated dark-noise is *<* 100 counts/sec for all detector elements. The scaling factor *w*_*i,j*_ are obtained normalising the signal registered by each element of the detector array after the subtraction of the dark noise.

As PSFs we used a simple Gaussian model which considers the shift-vector calculated *via* the phase correlation method. The full-width at half-maximum (FWHM) of the PSFs is estimated using the Fourier-ring-correlation (FRC) analysis on the adaptive pixel-reassignement STED-ISM image [11]. In short, the resolution of the STED-ISM image is calculated using the FRC analysis, the same value is used as FWHM for all the PSFs. This protocol results in a sort of blind reconstruction, where no input from the user is requested.

## Supplementary Material

**Figure S1:**
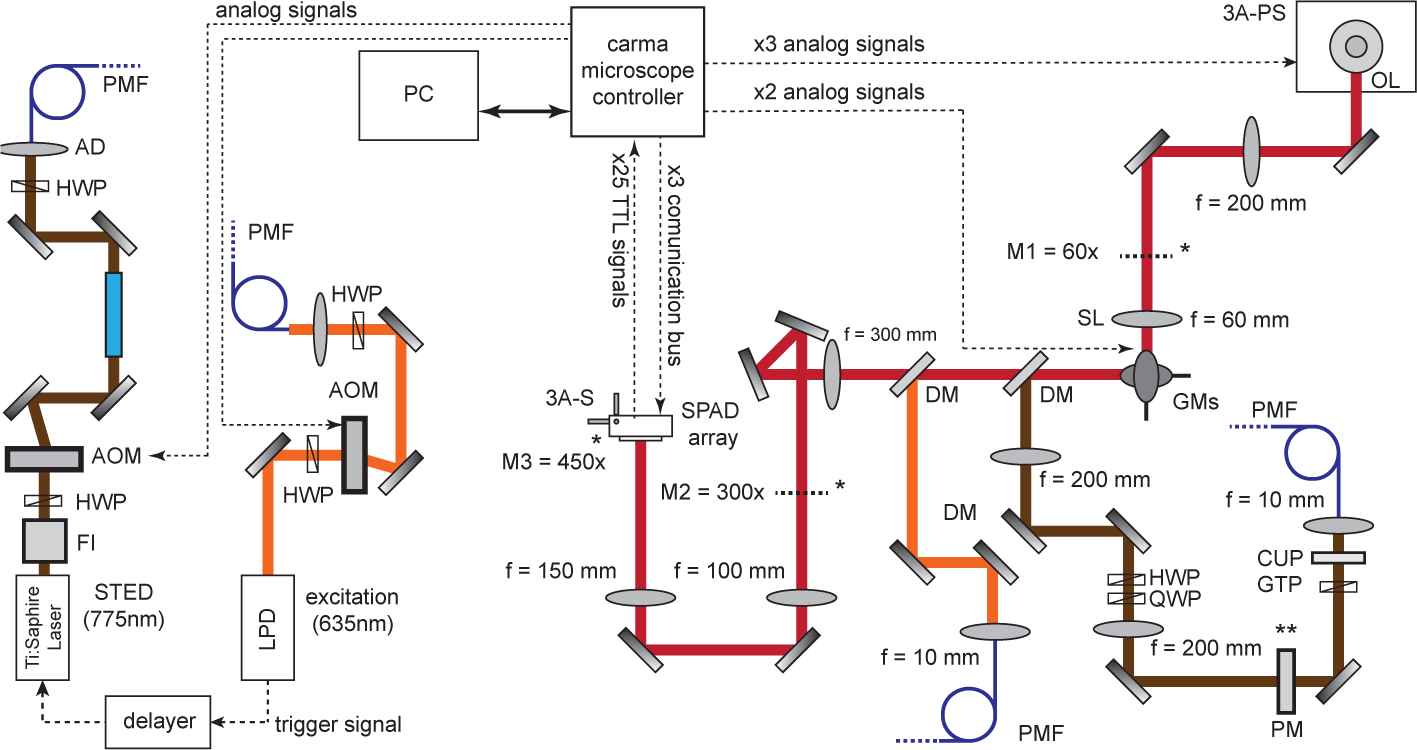
The STED-ISM setup. HWP:half-waveplate;AOM:acousto-optic modulator; AD:achromatic doublet; PMF: polarized-maintaining fiber; FI: Faraday isolator; GR: glass-road; 3A-PS: three-axis piezo stage; SL: scanning lens; GNs: galvanometric mirrors; DM: dichroic mirror; QWP: quarter-wave plate; BPF: band pass filter; MMF: multi-modes fiber; APD: avalanche photo-diode; PM: phase-mask; GTP: Glan-Thompson polarizer; CUP: clean-up filter. The asterisk denotes the plane conjugate to the image plane. The double asterisks denote the plane conjugate to the objective back-aperture.

**Figure S2:**
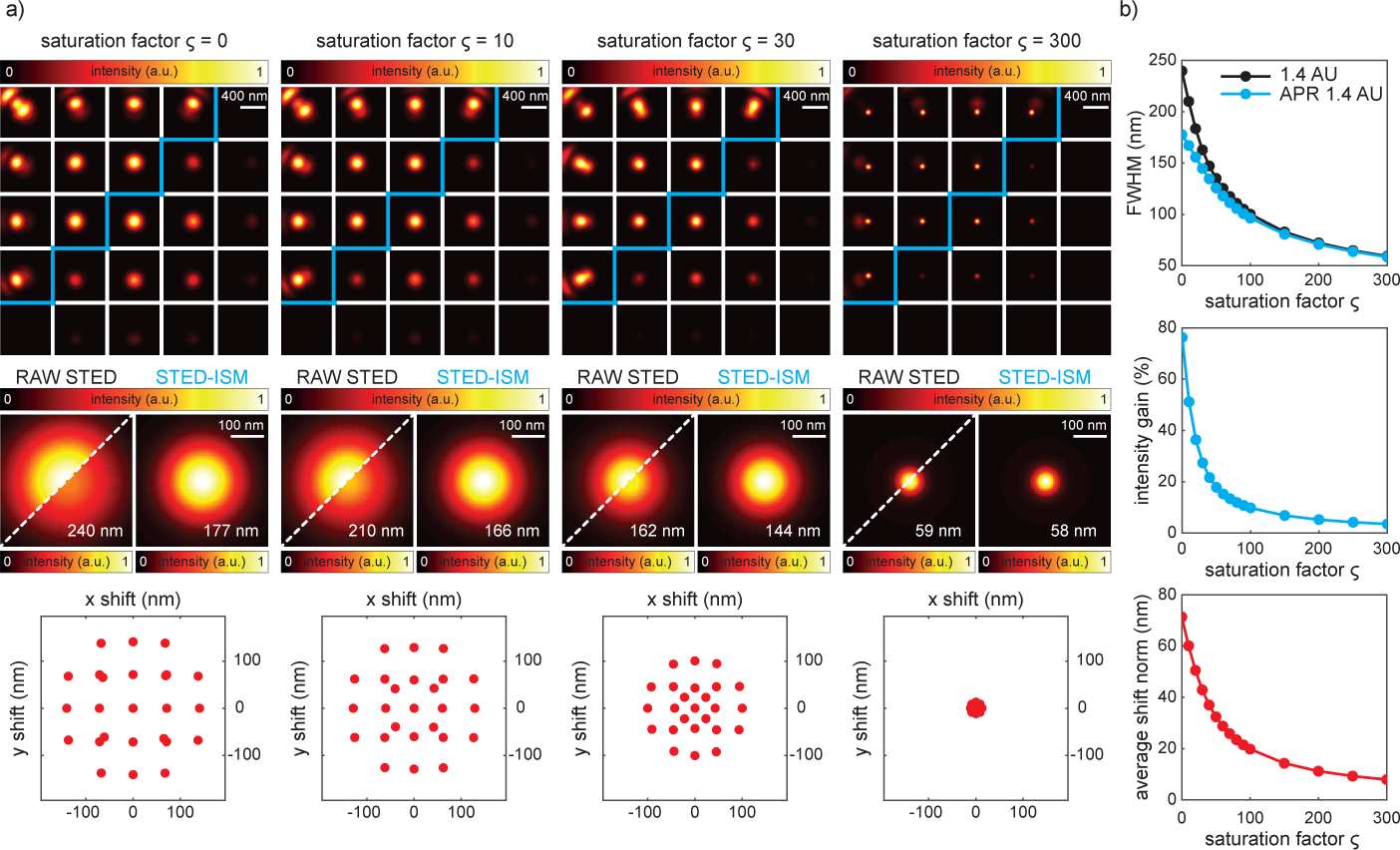
Simulation of a STED-ISM experiment with increasing saturation factor *ζ*. **a**, Each element of the SPAD array is characterised by a PSF which is, in first approximation, shifted with respect to the central one. The 25 PSFs are represented in the top row, normalized to themselves (top left), or to the central one (bottom right). Each PSF is shifted back, accordingly to the values retrieved by our phase correlation algorithm (bottom row), and the summed up to obtain the STED-ISM PSF (central row, right). The raw STED PSF is alternatively obtained by ignoring the shifts and only summing all the PSFs up (central row, left, normalized and non-normalized versions reported). FWHM values are also reported. As the saturation factor *ζ* increases (from left to right) and consequently the effective excitation PSF shrinks, the shift values reduce and the benefits of STED-ISM with respect to the raw STED counterpart become less and less evident. **b**, STED-ISM grants a resolution improvement (top, FWHM analysis) as well as a SNR increase (center) if *ζ* is lower than a given value. Bottom, the average of the four shortest shift values is decreasing as a function of *ζ*. Excitation, emission and STED wavelengths: 640, 670 and 775 nm respectively; STED repetition rate: 80 MHz; STED pulse length: 200 ps; SPAD array semi-length: 1.4 Airy Units.

**Figure S3:**
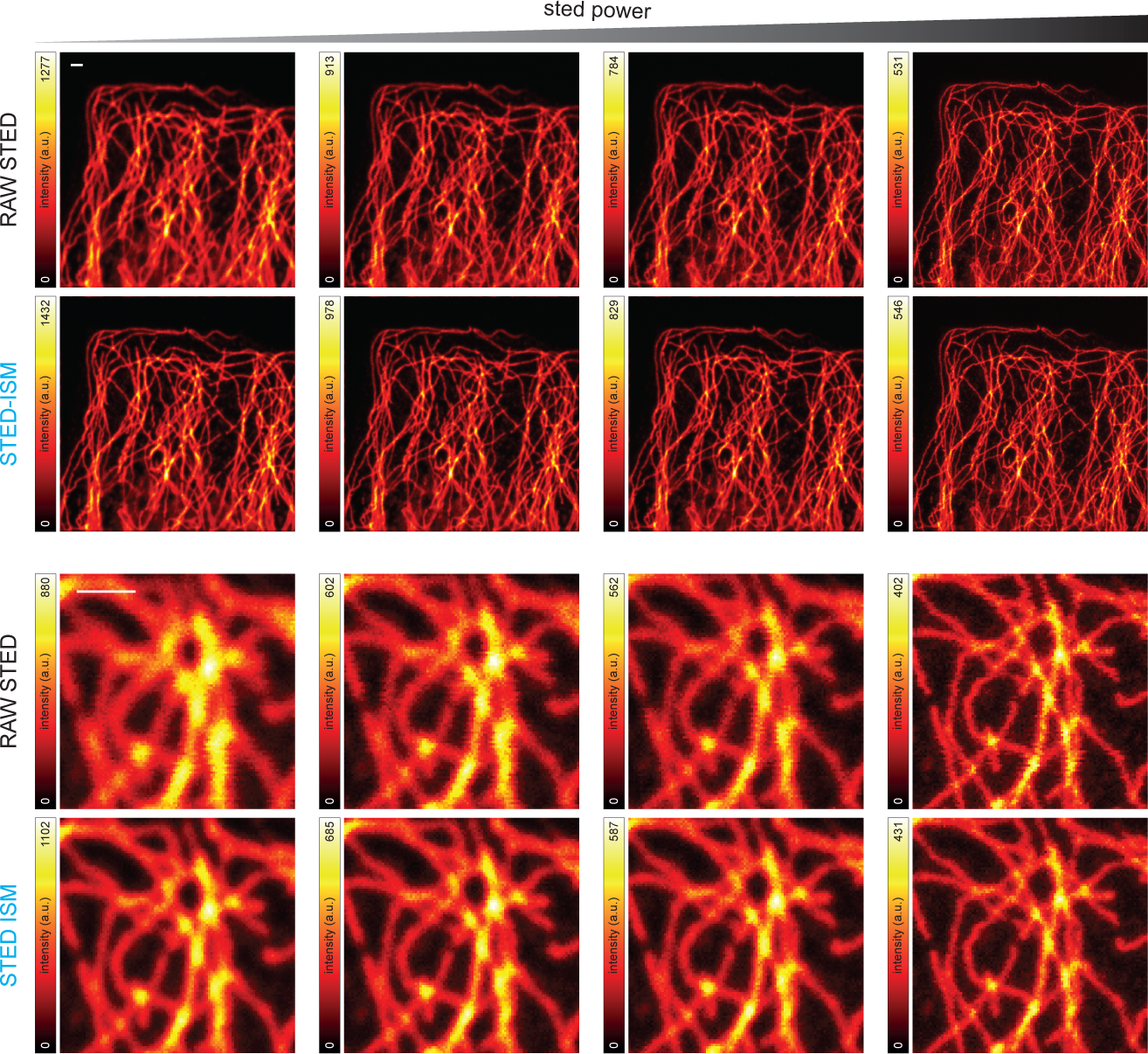
STED-ISM measurements of fixed cells with increasing STED power. The resolution and the SNR of are enhanced by STED-ISM when imaging tubulin-labelled fixed Hela cells. Retrieved FRC resolution values: 278 and 189 nm, 202 and 180 nm, 192 and 177 nm, 136 and 136 nm, for raw STED and STED-ISM, respectively. Dwell time: 100 *µ*s; pixel size: 40 nm; STED powers: 0, 15, 23, 137 mW; format: 500 × 500 pixels; details format: 150 × 150 pixels; all scale-bars: 1 *µ*m.

**Figure S4:**
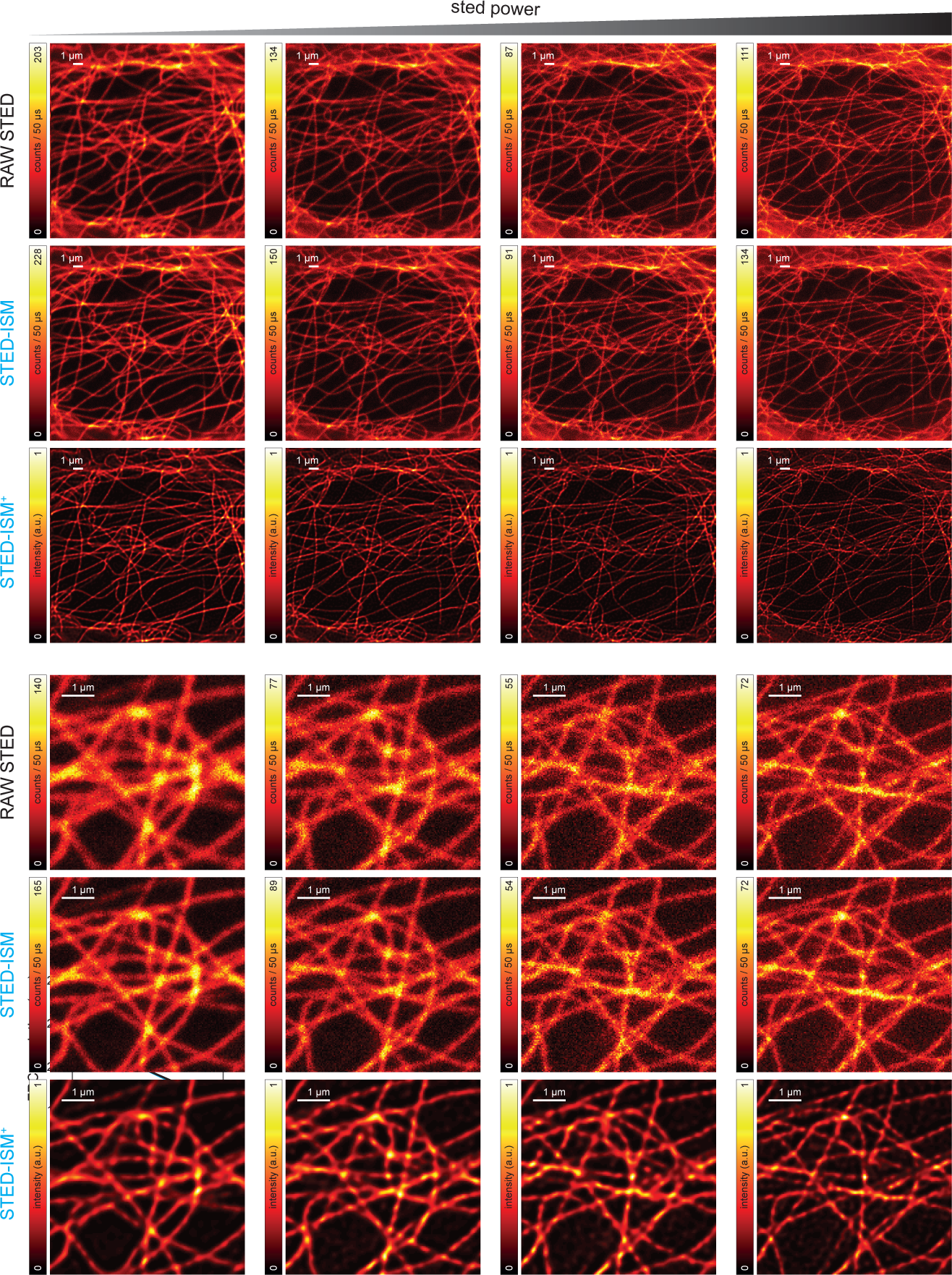
Live cell STED-ISM imaging with increasing STED power. Retrieved FRC resolution values: 290 and 213 nm, 233 and 202 nm, 202 and 202 nm, 168 and 165 nm, for raw STED and STED-ISM, respectively. Dwell time: 50 *µ*s; pixel size: 40 nm; STED powers: 0, 10, 20, 45 mW; format: format: 500 × 500 pixels ;details format: 150 × 150 pixels.

**Figure S5:**
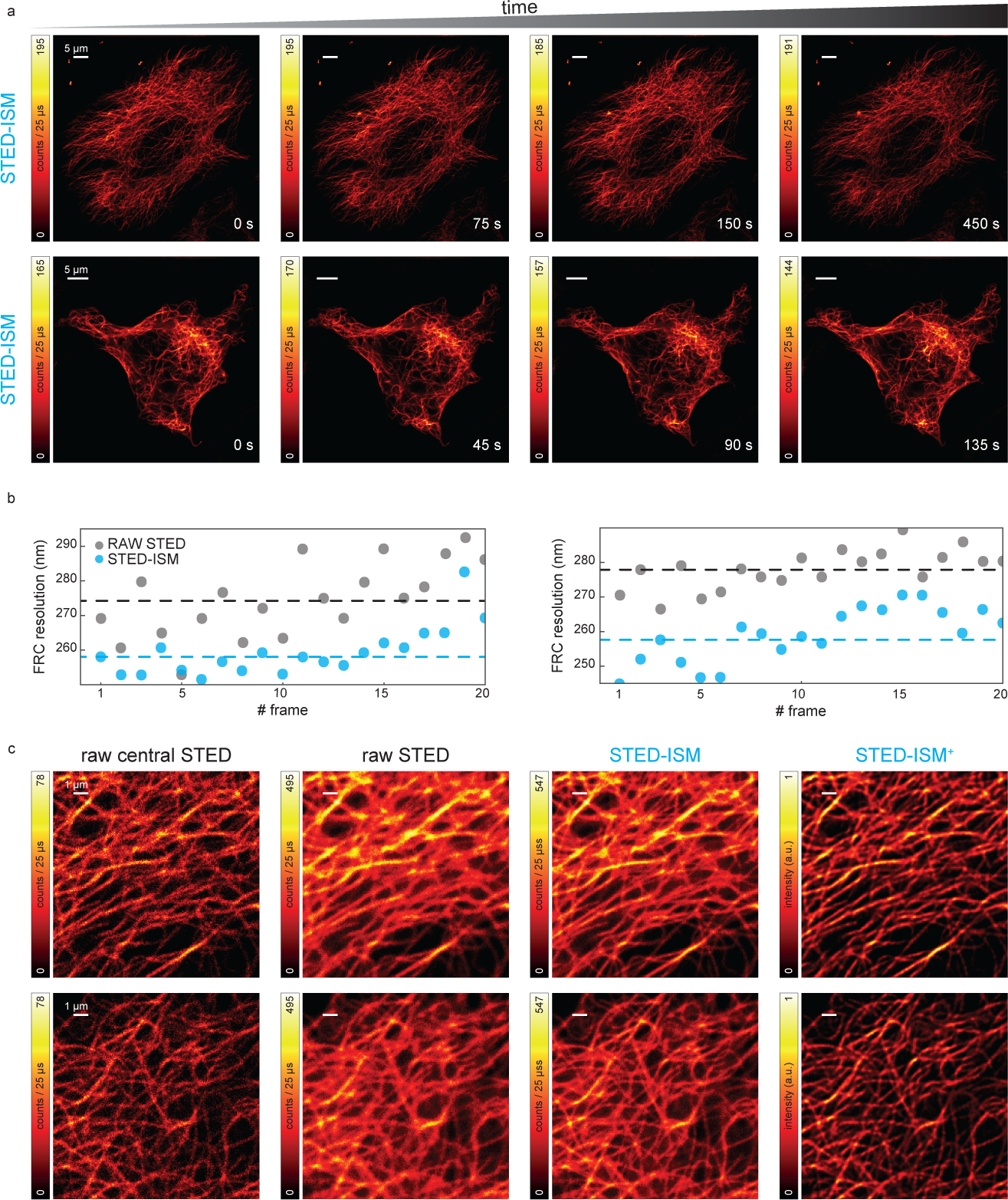
STED-ISM time lapses. **a**, Two extended time lapses (20 frames, 500 and 300 s) of Sir-Tubulin labelled Hela cells. Different frames are reported to show the negligible photo-bleaching. **b**, The FRC analysis confirms, for both time-lapses, the superior performances of STED-ISM with respect to raw STED. **c**, Details from the time lapses, showing the image recorded by the central element of the SPAD array (raw central STED), the result of the summing operation (raw STED), the result of the pixel reassignment operation (STED-ISM) and the deconvolution image (STED-ISM^+^). Dwell time: 25 *µ*s; pixel size: 70 and 67 nm; format size: 1000 × 1000 and 750 × 750 pixels; details format size: 200 × 200 pixels; 635 nm excitation laser power: 8 W; 775 nm STED laser power: 21 mW.

